# Antimicrobial resistance as a signature of soil restoration across a 143-year chronosequence

**DOI:** 10.64898/2026.06.02.729492

**Authors:** Tim Goodall, Briony Jones, Amy C. Thorpe, Robert I. Griffiths, Hyun Soon Gweon, Daniel S. Read, Richard Pywell, Susheel Bhanu Busi

## Abstract

Restoring agriculturally degraded habitats to species-rich grasslands is a vital conservation objective. During restoration, how the soil resistome matures alongside microbial community composition and function remains unclear. Here, we tested two competing hypotheses: whether the soil resistome matures through a microbial warfare model, in which restoration fosters higher-order biotic interactions, or whether antimicrobial resistance (AMR) is instead driven by the competitive pressures of the high taxonomic richness found in disturbed, eutrophic arable land. Using a unique land-use chronosequence on Salisbury Plain, UK, we investigated the trajectory of ecosystem reassembly following the cessation of agricultural activity. Our results demonstrate that AMR abundance increases significantly with restoration age, reaching a maximum in >143-year-old soils. Aligned with this rise in AMR abundance were significant increases in microbial biomass, dominance and cross-kingdom interaction as the ecosystem matured. This suggests that resistome expansion is not associated with generalised bacterial competition, but by a structural maturation of the microbiome. We observed an order of magnitude increase in biosynthetic potential, dominated by the emergence of streptomycin clusters. This maturation was characterised by a loss of bacterial diversity and a systematic shift towards high eukaryote-to-prokaryote ratios. This reorientation mirrors the expansion of a core resistome comprised of ancient, intrinsic mechanisms, such as MFS efflux pumps and RbpA target protection, in older soils. We demonstrate that endogenous AMR is a hallmark of healthy, restored soil ecosystems rather than a marker of anthropogenic degradation, positioning the resistome as bio-indicator of edaphic restoration success within calcareous soils.

## Main

The recovery of plant communities during grassland restoration is comparatively well described^1,2,3^, but the maturation of the below-ground microbiome and it’s resistome during soil recovery remains much less clear. In particular, there is a lack of certainty as to whether the soil resistome reflects disturbance^4^, competition^5^, and nutrient enrichment^6^ in arable systems, or instead emerges as a feature of long-term ecological recovery^7^. This distinction matters because AMR in soils is not solely a product of anthropogenic contamination^8^ but is also rooted in ancient microbial interactions^9^ and natural secondary metabolism^10^. Following our recent characterisation of chronological soil functional maturation^11^ using a multi-generational calcareous grassland restoration chronosequence on Salisbury Plain, UK, we here test how soil resistome composition and abundance change as soils mature from arable land, through restoration, to ancient grassland^12^. Grasslands were classified as arable (0 years; mean organic matter 10% LOI), early regenerating (27 years; 16% LOI), regenerating (63 years; 20% LOI), and ancient (>143 years; 23% LOI) (Supp. Tab. 1). By integrating metagenomic AMR profiles with microbial diversity, microbial biomass markers, eukaryote-to-prokaryote ratio, and biosynthetic gene cluster potential (Supp. Fig. 1, Supp. Methods), we asked whether resistome expansion is associated with disturbed, species-rich arable soils or with the functional maturation of restored ecosystems.

### Microbial reorientation, proliferation, dominance, and AMR accumulation

Analysis of microbial biomass biomarkers, and of the metagenomic resistome^13-15^ revealed soil restoration age was positively associated with absolute microbial biomass, which expanded across the chronosequence (*R*^2^ = 0.41, *P* < 0.001; Fig. 1A). This systemic biomass proliferation was accompanied by a linear accrual of the soil antibiotic resistome. Total AMR gene abundance scaled positively, and significantly, with expanding total microbial biomass and fungal biomass (Fig. 1B-C), however, the strongest predictor of resistome expansion was a structural shifting away from bacterial hegemony: the Eukaryote:Bacteria ratio strongly and positively predicted total AMR abundance (*R*^2^ = 0.64, *P* < 0.001; Fig. 1D). Furthermore, resistome accumulation was heavily favoured in consolidated, less even communities, showing a clear positive relationship with Simpson’s dominance index (*R*^2^ = 0.50, *P* < 0.001; Fig. 1E). Together, these results demonstrate that long-term soil successional development drives a predictable reorientation of the soil microbiome, where biomass expansion, fungal-eukaryotic progression, and community dominance collectively culminate in enhanced antibiotic resistome accumulation. The pattern of increasing fungal prominence in older soils (143 years) may reflect broader changes in substrate use, decomposition pathways, and the chemical environment experienced by bacterial communities.

**Figure 1.**
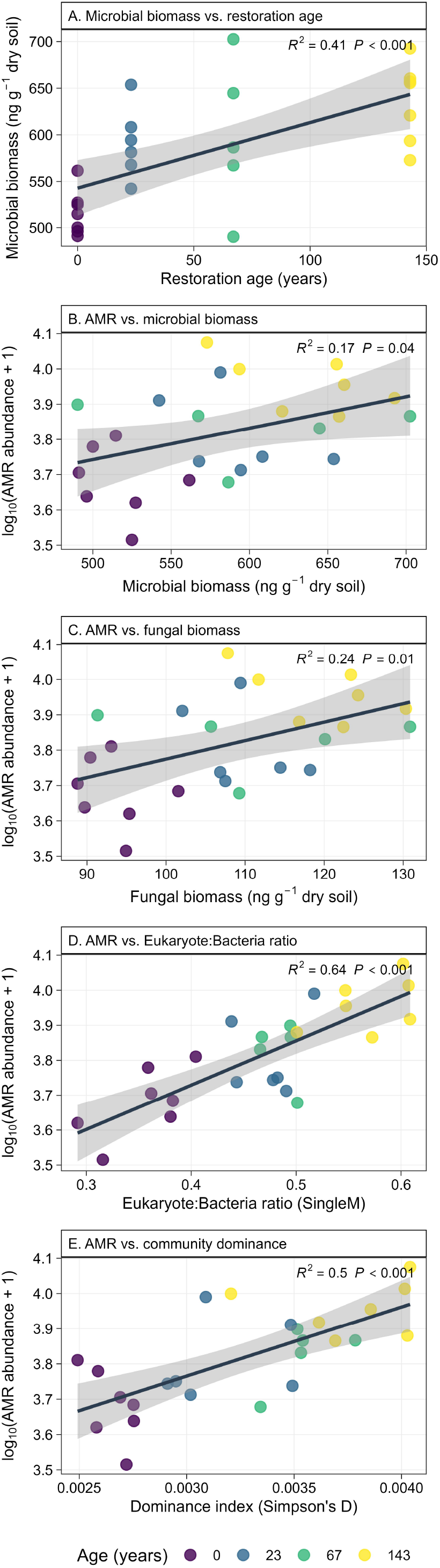
Ecological and biomass drivers of soil antibiotic resistome abundance across a restoration chronosequence. (A) Relationship between soil restoration age (years) and total microbial biomass (ng g^-1^ dry soil) derived from microbial phospholipid fatty acid (PLFA) profiles. (B) Total AMR gene abundance (log_10_ FPKPMC + 1) relative to total estimated microbial biomass. (C) Linear regression exploring the influence of absolute fungal biomass (ng g^-1^ dry soil) on total AMR abundance. (D) Direct relationship between the Eukaryote:Bacteria biomass ratio (SingleM marker-derived) and total AMR abundance. (E) Impact of community consolidation on the resistome, modeled as total AMR abundance against community dominance (Simpson’s D index calculated at the singleM OTU level). Individual data points represent independent soil core replicates, coloured by their respective chronosequence age. Solid lines indicate linear model fits calculated across the complete dataset, with shaded regions representing 95% confidence intervals. Evaluated goodness-of-fit (*R*^2^) and associated significance (*P* values) are annotated within each respective panel.

### Successional coupling of biosynthetic potential and natural resistance

The successional shift in the resistome was underpinned by a tenfold increase in biosynthetic potential. Total biosynthetic gene cluster (BGC) abundance increased by an order of magnitude from arable to ancient soils with unique gene clusters rising from a baseline of ∼5 in arable soils to >50 in ancient grasslands (Fig. 2A). This substantial increase in biosynthetic potential is consistent with a plausible ecological basis for the parallel rise in endogenous resistance genes because BGCs and resistance genes are typically co-selected^16^ (Fig. 2B; Supp. Fig. 2). Resistome analysis across the chronosequence revealed successional increases in efflux systems (MFS, RND, and ABC transporters) and target protection mechanisms (RbpA) which safeguard essential cellular machinery from natural secondary metabolites^17^. The enrichment of streptomycin-like BGCs in ancient soils suggests that mature systems harbour a more specialised secondary metabolome, likely reflecting intensified antagonism between prokaryotic and eukaryotic communities^16^.

**Figure 2.**
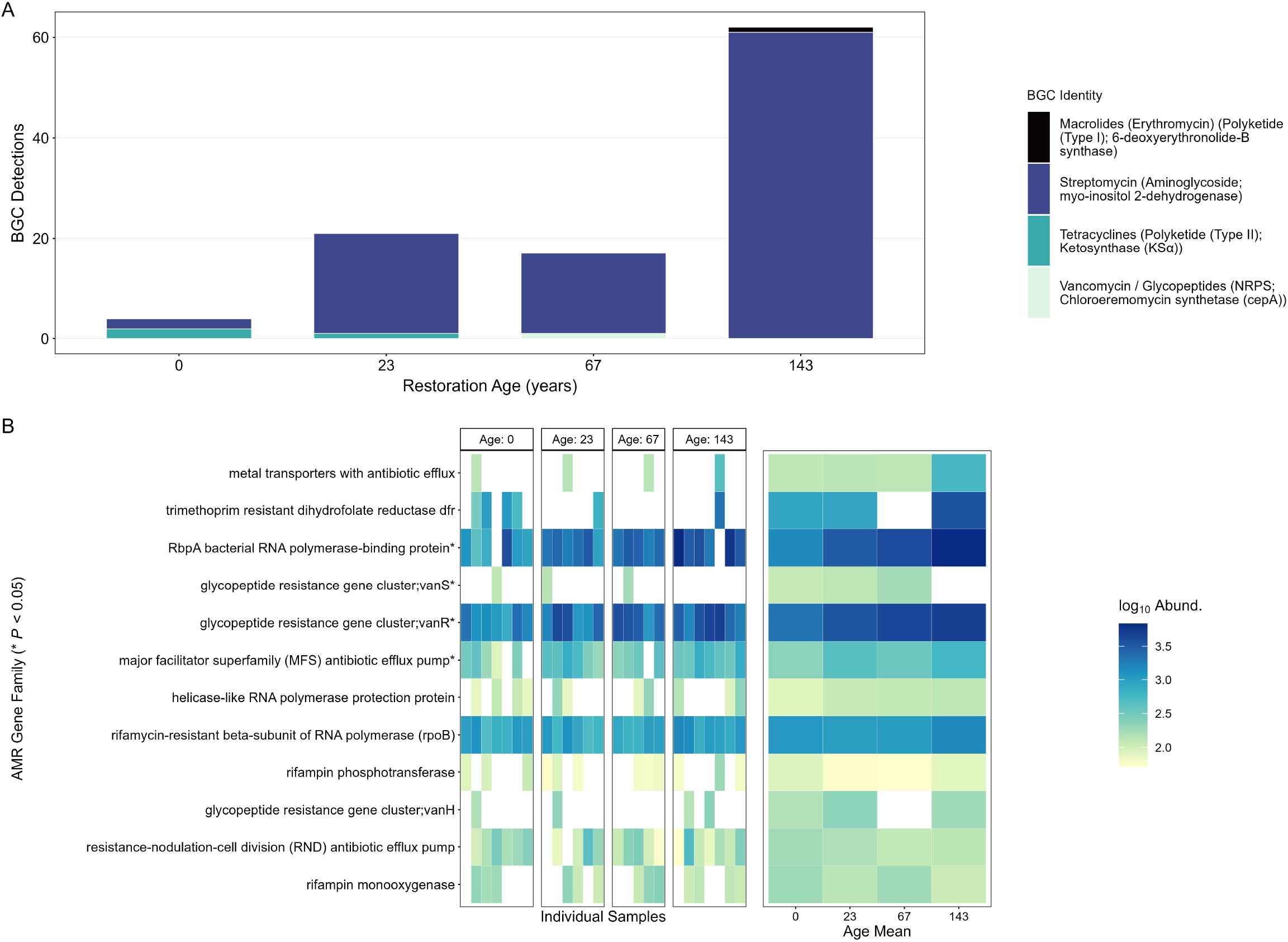
Succession of Soil Antibiotic Resistome and Biosynthetic Potential. (A) Quantitative occurrence of Biosynthetic Gene Clusters (BGCs) across successional stages. Categorical fill levels represent a hierarchical classification: Group (e.g., Macrolides), followed by Classification (e.g., Polyketide Type I), and predicted Enzymatic Function (e.g., 6-deoxyerythronolide-B synthase). Total detections represent the sum of all “Bio” type clusters identified within each age group. (B) Heatmap illustrating the log_10_ abundance (FPKPMC) of AMR gene families across the restoration chronosequence. Data are presented for individual soil samples faceted by restoration age (left), with the corresponding “Age Mean” (right) highlighting age-level shifts. Gene families marked with an asterisk (*) represent significant linear responses to restoration age (*P* < 0.05).

### Conclusion

Our findings challenge the assumption that environmental AMR reflects anthropogenic degradation^8^. Across this 143-year chronosequence reflecting natural regeneration of calcareous grassland, resistome abundance increased alongside broader shifts in microbiome structure and biosynthetic potential, indicating that endogenous AMR can also be a feature of long-term ecological recovery. These results highlight the importance of distinguishing natural, restoration-associated resistome signals from those linked to anthropogenic AMR pressure. We observed that clinical markers like rifampin monooxygenase remained static, while natural markers linked to soil BGCs were strongly enriched in mature sites, suggesting that AMR composition and abundance may provide a useful high-resolution indicator of microbiome reassembly and restoration status in calcareous grassland soils.

## Notes

### Competing Interest Statement

The authors have declared no competing interest.

### Funding

- Natural Environment Research Council: NE/S005137/1
- Biotechnology and Biological Sciences Research Council: BBX011089/1

