## Supplementary material for "Antimicrobial resistance as a signature of soil restoration across a 143-year chronosequence": Methods

***Supplementary Methods.***

*Study site, sampling and environmental characterisation*

Salisbury Plain, managed by the UK Ministry of Defence, covers approximately 38,000 ha and contains Western Europe’s largest remaining area of lowland calcareous grassland. Its long-term use as a military training area has limited agricultural intensification, creating a landscape suitable for examining grassland restoration chronosequences. Site selection followed the land-history classifications of Redhead^1^, derived from historic map series. In total, 25 sites were sampled: seven arable, six recent grassland, five middle-aged grassland and seven ancient grassland sites. These represented minimum unimproved grassland ages of 0 years for arable land, 23 years for recent grassland, 67 years for middle-aged grassland and 143 years for ancient grassland.

Soils were collected in January 2013. At each site, five subsamples were taken along a 100 m transect using 5 cm plastic corers to a depth of 15 cm, or until an impenetrable chalk horizon was reached. Soil below the organic horizon was homogenised, transported to the laboratory and stored at −20 °C.

Vegetation surveys were conducted in summer 2013 along the same transects. Vascular plant cover was recorded in five 2 × 2 m quadrats placed at 20 m intervals and scored using the DAFOR scale. For Shannon diversity calculations, DAFOR scores were converted as follows: Dominant = 5, Abundant = 4, Frequent = 3, Occasional = 2, Rare = 1 and Present = 0.1.

Soil physicochemical analyses included pH, total and organic carbon, phosphorus, potassium, magnesium and texture, measured by NRM-Cawood Scientific. Soil moisture, organic matter by loss on ignition, total nitrogen and organic nitrogen were measured at the UKCEH Centralised Chemistry laboratories using established protocols. C:N ratios were calculated from organic carbon and organic nitrogen values.

*AMR profiling*

To profile the soil resistome, metagenomic reads were functionally annotated using a dual-pipeline approach. Specialised antimicrobial resistance genes (ARGs) were identified by querying non-redundant gene catalogues using ResScan (<https://github.com/hsgweon/resscan>), with the CARD database^2^. Simultaneously, broad functional categorisation was performed using eggNOG, mapping sequences to the eggNOG orthology database to capture intrinsic resistance determinants and associated metabolic pathways. To account for differences in sequencing depth and gene length, abundance was normalised to Fragments Per Kilobase per Million mapped reads onto the Chromosome (FPKPMC). This normalization provides a robust proxy for gene copy number per microbial cell by using the abundance of single-copy marker genes as a genomic denominator, allowing for direct quantitative comparisons of the resistome across the restoration chronosequence.

*Eukaryote-to-prokaryote ratio*

To quantify the relative abundance of eukaryotic and prokaryotic communities, we calculated the eukaryote-to-prokaryote ratio using domain-specific marker gene abundances from the SingleM pipeline^3^. SingleM utilizes a suite of 14 universal, single-copy ribosomal protein marker genes (e.g., rplB, rpsC) to estimate the relative cellular abundance of Bacteria, Archaea, and Eukaryotes. By using single-copy markers, this approach provides a robust proxy for the ratio of eukaryotic to prokaryotic cells, bypassing the biases typically associated with varying genome sizes and 16S/18S rRNA gene copy number variations across different microbial taxa.

*Microbial diversity*

SingleM marker-gene-based abundance estimates were used to produce SingleM abundance tables. For each sample alpha diversity values for microbial richness, Shannon diversity, and Simpson dominance were generated based on the counts obtained via SingleM.

*Microbial biomass*

To resolve absolute microbial community biomass, total lipid extracts obtained from fresh soil samples were structurally analysed using Gas Chromatography (GC)^4^. Raw peak profiles were filtered to isolate soil microbial signalling from confounding macro-organisms and plant inputs. High-order plant waxes and macrofauna structural lipids—specifically C20:0, C22:0, C24:0, C18:3ω6c, C18:3ω3c, C20:4ω6c, and C20:5ω3c—were excluded from baseline community metrics. The cumulative mass of all remaining active microbial phospholipid fatty acid (PLFA) fragments was summed to yield the total microbial lipid yield. Fungal markers - C18:1ω9c, C18:2ω6c, C18:2ω9t. Bacterial markers (Gram+ branched, Gram- monoenoics, cyclopropyl, actinomycetes) – “i-C15:0”, “a-C15:0”, “i-C:16:0”, “i-C17:0”, “a-C17:0”, “i-C18:0”, “a-C18:0”, “C16:1ω7c”, "C16:1ω9c", "C17:1ω10c", "C18:1ω11c", "C18:1ω11t", “cy-C17:0*”, “9,10-cy-C19:0”, “11,12-cy-C19:0”, "10Me-C16:0", "10Me-C17:0*", "10Me-C18:0*". Gravimentric measurements of soil moisture were used to correct the microbial yield to ng g^-1^ dry soil.

*Biosynthetic gene clusters*

Biosynthetic Gene Clusters (BGCs) were identified and characterised using an integrated genomic mining pipeline. Metagenomic assemblies were processed through antiSMASH v7.0 (Antibiotic and Secondary Metabolite Analysis Shell)^5^ and GECCO (Gene Cluster Prediction with Conditional Random Fields)^6^ to ensure a comprehensive capture of both canonical and novel secondary metabolic pathways. To provide high-resolution functional annotation, predicted clusters were cross-referenced against the MIBiG (Manual of Minimum Information about a Biosynthetic Gene)^7^ database. A cluster was considered a 'known match' if it exhibited high homology to experimentally characterized BGCs (e.g., >40% gene similarity). Total BGC potential was quantified as the non-redundant sum of clusters identified across all platforms, normalised to the total number of contigs >5kb to account for assembly fragmentation. Redundant BGC predictions across antiSMASH and GECCO were consolidated based on genomic coordinates, ensuring each locus was counted only once in the final quantitative analysis.

*Data and code availability*

Sequence data is available at NCBI-SRA under BioProject ID: PRJNA1424699. Scripts and data tables are available at https://doi.org/10.5281/zenodo.20396655
