## Supplementary figures and images for "Antimicrobial resistance as a signature of soil restoration across a 143-year chronosequence"

### Supp Fig 1

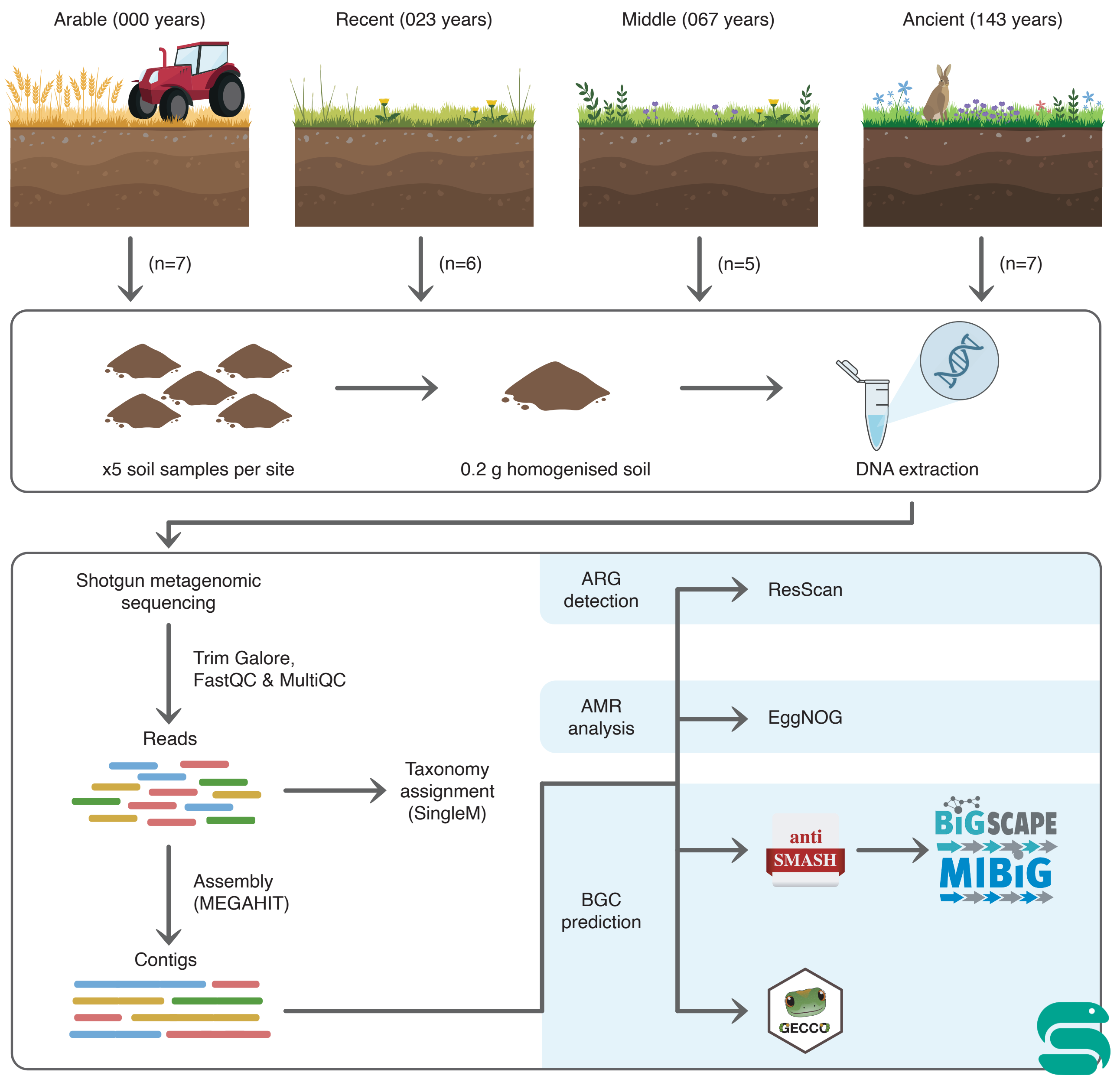

### Supp Fig 2

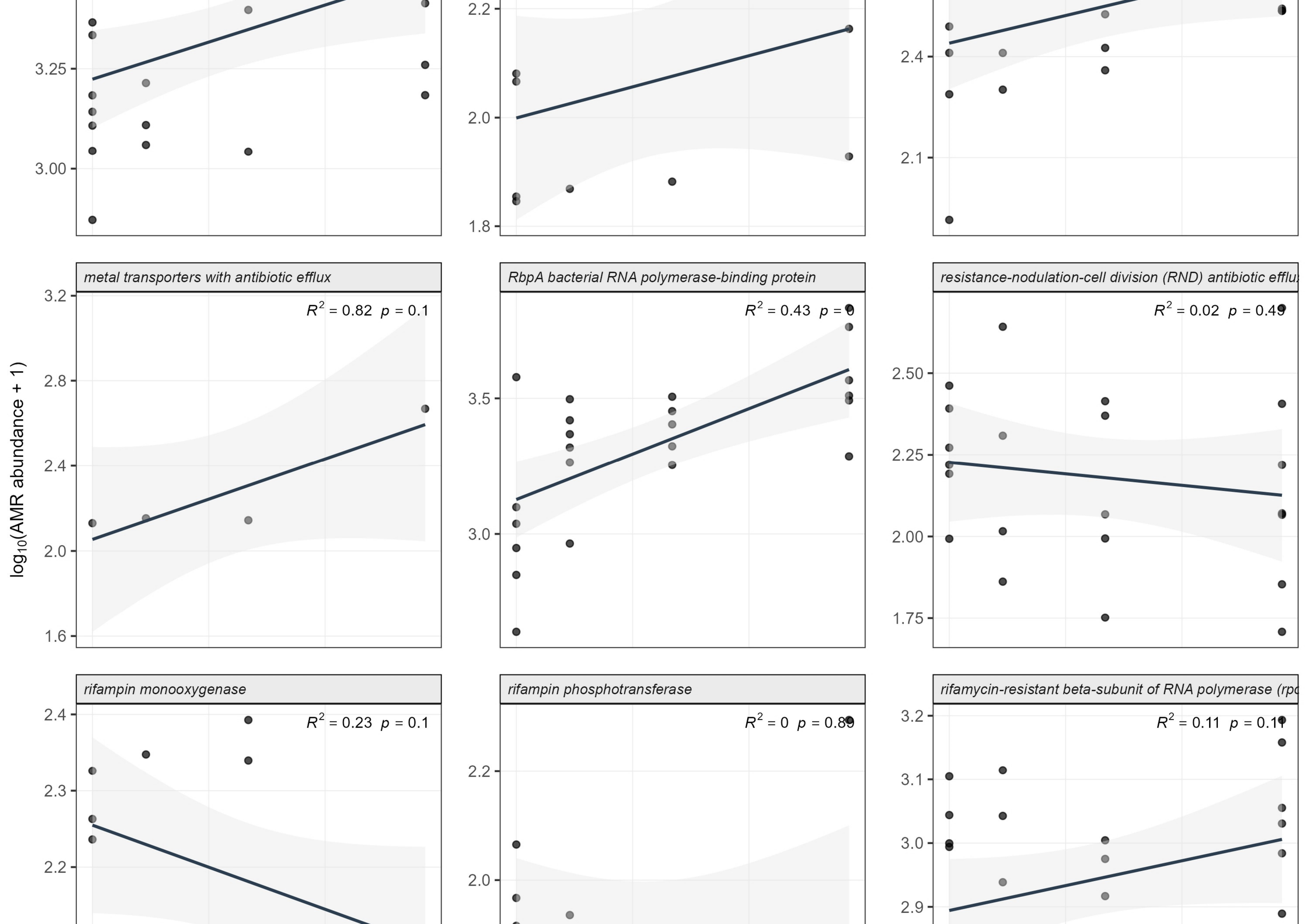
