## Supplementary material for "Antimicrobial resistance as a signature of soil restoration across a 143-year chronosequence": Supp Legends

**Supplementary Figure 1. From Field to Findings.** A schematic of the sample analysis workflow. Full details of sites, sample processing, sequence data generation, tools and analysis are available in Supplementary Methods section.

**Supplementary Figure 2. Linear Regression Analysis of Core AMR Gene Family Responses.** The successional trends of nine prevalent Antimicrobial Resistance (AMR) gene families. Each panel represents a specific gene family, as identified by Homolox, with log_10_-transformed abundance (FPKPMC) plotted against soil restoration age (years). Solid lines indicate the linear model (LM) fit, shaded regions represent the 95% confidence interval. Statistical annotations within each panel (*R*^2^ and *P* value) were calculated using a standard linear regression model. Points represent individual soil samples. Families are faceted with independent y-axis scales to visualise relative shifts in abundance across the chronosequence.

**Supplementary Table 1. Measured soil variables.** Mean values across the chronosequence.
